# Crowding distorts features, not objects: structured target confusions in a 4AFC conjunction task

**DOI:** 10.64898/2026.10.06.756933

**Authors:** John Cass, Erik Van der Burg

## Abstract

Visual crowding is the disruption of peripheral object identification by nearby flanking clutter. It remains unclear whether crowding disrupts the bound representation of an object or degrades the object’s component features separately. We addressed this with a 4AFC conjunction-report task in which observers reported both the orientation and the brightness of every target. Target Gabors at 6° eccentricity were presented alone or flanked at 1, 3, or 5 degrees separation by Gabors varying independently in colour and/or orientation similarity to the target. We analysed the data three ways: joint accuracy, accuracy on each feature alone, and the statistical coupling between the two report channels. Orientation reports were principally impaired by orientation-similar flankers within the smallest target–flanker separations, whereas luminance reports were impaired by target-colour-similar flankers across a broader spatial range. Observed joint accuracy was closely predicted by the product of the two marginal accuracies in every condition. A complementary chi-square analysis of the full 4x4 stimulus-response matrices agreed with this conclusion at the most-crowded conditions, while revealing structured response biases that the ratio test could not detect. These biases were present at baseline and were not amplified by crowding. The conjunction errors observed under crowding therefore arose from two largely independent feature-specific failures, rather than from disruption of a representation in which the two features are bound. These results indicate that object-level crowding errors in this paradigm are the downstream signature of two independent feature-level processes rather than the breakdown of an object-level mechanism.

## Introduction

Peripherally presented objects are often more difficult to identify when located in cluttered visual environments than when presented in isolation, a phenomenon known as visual crowding (Whitney & Levi, 2011). Two stimulus-based principles are well established as determinants of crowding strength. First, crowding tends to be stronger when target and nearby flanking elements share similar features (Kooi et al., 1994). Second, and more fundamentally for its measurement, crowding depends on target–flanker proximity: impairment is greatest when flankers fall within a critical spacing region around the target, roughly half the target’s eccentricity in radius, a relationship known as Bouma’s law (Bouma, 1970). For this reason, crowding is typically quantified as a reduction in target-identification accuracy as flankers approach the target within this critical spacing window (Pelli & Tillman, 2008).

Accuracy, however, tells us only *that* the observer was wrong. It does not reveal *how* the percept failed. In a standard binary forced-choice task, an incorrect response could reflect a non-specific loss of target information, substitution of the target by a flanker (Dakin et al., 2010; Harrison & Bex, 2017; Strasburger & Malania, 2013), compulsory averaging of target and flanker features (Dakin et al., 2010; Greenwood et al., 2009; Parkes et al., 2001), uncertainty about one particular feature dimension, and/or failure to bind multiple target features into a coherent object identity (Pelli et al., 2004; Poder & Wagemans, 2007; Treisman & Schmidt, 1982). These possibilities have distinct theoretical implications, yet all collapse into the same behavioural outcome: an incorrect response.

A multi-alternative forced-choice task provides a way to open up this error structure. Dakin et al. (2010) demonstrated this principle using a 4AFC paradigm in which observers reported the global orientation of a letter-like T target, i.e. one of four possible target orientation states (T, ⊥, ⊣, ⊢) defined by the conjunction of the positions of the target’s component features. Rather than focusing on overall error rates, they examined the full pattern of target–response confusions trial by trial. This approach revealed that crowding errors are not random: errors induced by end-flankers closely resembled the flanking stimulus, whereas errors induced by side-flankers were far less structured. Critically, a feature-level positional averaging model could account for these apparently object-level effects, demonstrating that structured multi-alternative stimulus-response matrices place stronger constraints on models of crowding than accuracy measures alone.

The present study extends this stimulus-response matrix approach to a conjunction design in which the target varies independently in two separate feature dimensions (orientation and brightness), producing four possible target identities. Because these two dimensions are orthogonally and independently varied, every response can be decomposed into a correct conjunction report, an orientation-only error (correct brightness, incorrect orientation), a brightness-only error (correct orientation, incorrect brightness), or a both-feature error (both dimensions incorrect). This decomposition is not available in single-feature tasks or in tasks where the response alternatives are defined by a single integrated property. Column-normalised 4×4 stimulus-response matrices therefore reveal not only whether crowding occurred, but whether the resulting distortion was specific to one feature dimension or affected the target conjunction as a whole.

Different accounts of crowding make different predictions for this two-feature error structure. If crowding reflects a non-specific degradation of target information, errors should be distributed relatively uniformly across all three incorrect response categories regardless of flanker type. If, by contrast, crowding operates primarily at the level of individual feature representations that are largely independent of one another (the *independent features hypothesis* (Cass & Van der Burg, 2023; Greenwood & Parsons, 2020; Yashar et al., 2019), then the pattern of errors should be contingent upon the specific feature relation between target and flankers. Colour-related flanker structure should selectively distort brightness reports, and orientation-related flanker structure should selectively distort orientation reports. Under this account, both-feature errors should be largely predictable from the independent co-occurrence of single-feature errors, because orientation and brightness are corrupted at separate processing stages before being combined into a response.

A stronger alternative, the *bound features hypothesis*, holds that crowding can disrupt the binding of features into a coherent object representation (Pelli & Tillman, 2008; Yashar et al., 2019). According to this account, conjunction errors should be super-additive: both-feature errors should occur more frequently than expected from the marginal rates of single-feature errors, particularly under the strongest crowding conditions.

In a previous study using this same stimulus set and flanker manipulations, we demonstrated a double dissociation between task and the feature dimension defining target–flanker similarity (Cass & Van der Burg, 2023). Brightness judgments were highly contingent upon target–flanker colour similarity and largely unaffected by flanker orientation, whereas orientation judgments were primarily contingent upon the relative orientation of target and flankers. This pattern supports the independent features hypothesis in the context of separate, blocked tasks. The present study extends this work by requiring subjects to report both orientation and brightness on every trial. This conjunction design allows us to ask whether the independence of orientation and colour crowding (previously inferred from blocked task comparisons, and from conjunction-error patterns in tasks where response alternatives were defined by the bound positions of object features (Dakin, et al. (2010)) also holds when the two defining dimensions are orientation and brightness varied independently, or whether reporting two orthogonal features simultaneously reveals an additional object-level crowding process not visible in single-feature tasks.

To this end, we employed a 4AFC conjunction version of the paradigm from Cass and Van der Burg (2023). Subjects reported the full orientation–brightness conjunction of a peripheral Gabor target on each trial. Flankers varied in target–flanker separation, colour similarity, and orientation similarity. We first established whether conjunction accuracy showed the standard separation-dependent crowding profile, then used column-normalised stimulus-response matrices to determine whether the resulting errors were uniform or feature-contingent. We also tested whether both-feature errors exceeded predictions based on the independent co-occurrence of single-feature errors.

By moving from binary accuracy to the full structure of the stimulus-response matrix, the present study asks whether visual crowding disrupts object identity as a whole, or whether apparent object-level failures are built from separable distortions of component features.

## Method

### Participants

Thirty-nine subjects participated in the experiment, none of whom participated in our earlier study (Cass & Van der Burg, 2023). All had normal or corrected-to-normal vision. Informed consent was obtained prior to participation. The research was conducted in accordance with the ethical guidelines of the Faculty of Behavioural and Movement Sciences at the Vrije Universiteit Amsterdam and with those laid down in the Declaration of Helsinki.

Following the convention of our earlier work (Cass & Van der Burg, 2023), data from two subjects were excluded prior to analysis because their performance on target-alone trials was at or below chance (4AFC chance = 0.25; one subject 0.09, one subject 0.24). The remaining 37 subjects were retained for analysis (mean target-alone conjunction accuracy = 0.73, range 0.34–0.95).

### Apparatus and stimuli

The experiment was run in a dimly lit cubicle and programmed using OpenSesame software (Mathot et al., 2012). Subjects used a chinrest located 57 cm from an LCD monitor (120 Hz refresh rate). The stimulus display was presented on a black background (< 0.5 cd/m²).

On each experimental trial, three small red dots (27 cd/m²) appeared on screen for 500 ms: one fixation dot at the centre of the display, and two location markers at 6° to the left and right of fixation signalling the possible target locations. A green target Gabor stimulus then appeared for 150 ms at one of the two peripheral locations (randomised across trials). This brief presentation interval was used to discourage shifts in fixation toward the target location.

Each target Gabor had a sinusoidal luminance profile recruiting only the green component of the RGB display, with a spatial frequency of 4 cycles per degree of visual angle (d.v.a.) and a Gaussian envelope (σ = 0.25°). The green luminance modulation profile of each Gabor ranged from a trough of < 0.5 cd/m² to a peak of either 48 cd/m² (RGB 0, 255, 0; “bright” targets, contrast = 95) or 14 cd/m² (RGB 0, 145, 0; “dim” targets, contrast = 27), where contrast is defined as (Lₘₐₓ − Lₘᵢₙ) / Lₘᵢₙ. The orientation of each Gabor was either +7° or −7°from vertical. Target brightness and orientation were each selected randomly from trial to trial, as was target location (left or right visual field).

On flanker-present trials, two equidistant Gabor flankers were presented above and below the target. Flankers varied in colour and orientation to produce four feature combinations defined by crossing colour similarity to target (similar: green, RGB 0, 200, 0, 31 cd/m², contrast = 61; dissimilar: red, RGB 200, 0, 0, 14 cd/m², contrast = 27) with orientation similarity to target (similar: vertical, 0°; dissimilar: horizontal, 90°). Target–flanker centre-to-centre separation was 1 d.v.a. (near, within Bouma’s critical spacing window for our 6° eccentricity), 3 d.v.a. (intermediate), or 5 d.v.a. (far, outside Bouma’s window). Target-alone trials used no flankers.

### Task and procedure

Subjects reported the full conjunction of the target’s orientation and brightness on every trial using a 4AFC keyboard response. The four response keys (1, 2, 4, 5 on a standard keyboard) mapped onto the four possible target identities as follows: 1 = dim + left-tilt; 2 = dim + right-tilt; 4 = bright + left-tilt; 5 = bright + right-tilt. Target location (left or right visual field) varied across trials but was not relevant to the response. Subjects were instructed to maintain fixation on the central red dot throughout the experiment and were told that if they were unsure about the correct response they should nonetheless make a response.

All trial factors (target location, target brightness, target orientation, flanker colour, flanker orientation, and target–flanker separation) were selected randomly on each trial. Each subject completed eight blocks of 64 trials (512 trials total), separated by self-paced rest breaks, yielding approximately 128 target-alone trials and 32 trials per flanker condition at each separation level per subject.

### Data analysis

Analyses were conducted in R (v4.3.3; (Team, 2024)) using the lme4 package (v1.1-35.1; (Bates et al., 2015)). Practice trials were excluded. Each trial yielded two binary outcomes: whether the orientation response matched the target orientation, and whether the luminance response matched the target luminance. From these we derived three trial-level outcomes that we analysed in turn: *conjunction-correct* (both orientation and luminance correctly reported); *orientation-correct* (orientation correctly reported, regardless of luminance response); and *luminance-correct* (luminance correctly reported, regardless of orientation response).

Because each outcome was binary on a given trial, we modelled each with a generalised linear mixed-effects model (GLMM) with a binomial response distribution and logit link function. The GLMM is the analogue of repeated-measures analysis of variance for binary trial-level outcomes. Each model included target–flanker separation (1°, 3°, 5°), flanker colour similarity to target (similar, dissimilar), flanker orientation similarity to target (similar, dissimilar), and their two- and three-way interactions as fixed effects, with participant as a random intercept. Fixed effects were sum-coded so that the intercept corresponds to the unweighted grand mean of the cell means and so that interaction tests do not depend on which level of each factor is treated as the reference. The significance of each model term was assessed by a likelihood-ratio test, comparing the full model against a reduced model with that term removed. Where a significant interaction was decomposed, planned contrasts were conducted on the estimated marginal means (EMMs) with Holm correction applied within each family of comparisons. EMMs are reported on a probability scale. Effect-size differences between conditions are also reported in logit units.

We considered a maximal random-effects structure including random slopes for all main effects and two-way interactions (the three-way slope was inestimable given twelve cells per participant). The maximal structure produced singular fits with between-slope correlations at ±1, and in two of our outcomes also produced unstable likelihood-ratio statistics and a fixed-effect estimate that diverged to a near-boundary value while its standard error remained tightly constrained. An internally inconsistent solution that indicated the maximal structure was overfit relative to the information in our data. Following Bates et al. (2015) and Matuschek et al. (2017), we report results from a random-intercept-only model for all outcomes. This model converged cleanly in each case and yielded EMMs that tracked observed cell proportions to within 0.03 across all twelve cells of the design.

The target-alone reference values reported in the Results were computed from each participant’s proportion correct on target-alone trials, summarised across participants using the mean and between-subjects 95% confidence interval.

## Results

For all GLMM analyses, data from two participants were excluded because their performance was at or below chance in the target-alone condition (one near chance, one below chance – see *Participants* section above), matching the convention of our earlier work (Cass & Van der Burg, 2023). Practice trials were excluded. This left thirty-seven participants, with 14,208 flanked trials and 4,735 target-alone trials for analysis.

We analysed trial-wise responses on the 4AFC conjunction-report task at three levels of description. We first examined accuracy on the joint outcome: whether the participant correctly reported both target features. We then examined accuracy on each feature considered alone, to ask whether the joint result reflects two qualitatively distinct flanker effects. Finally, we directly tested whether the orientation- and luminance-report channels operate independently across the design. Target-alone reference values were computed from each participant’s proportion correct on target-alone trials, summarised across participants using the mean and 95% confidence interval based on the *t* distribution: conjunction accuracy = 0.708, 95% CI [0.649, 0.768]; orientation accuracy = 0.891, 95% CI [0.843, 0.939]; luminance accuracy = 0.788, 95% CI [0.749, 0.827]. Figure 1 summarises these findings.

**Figure 1.**
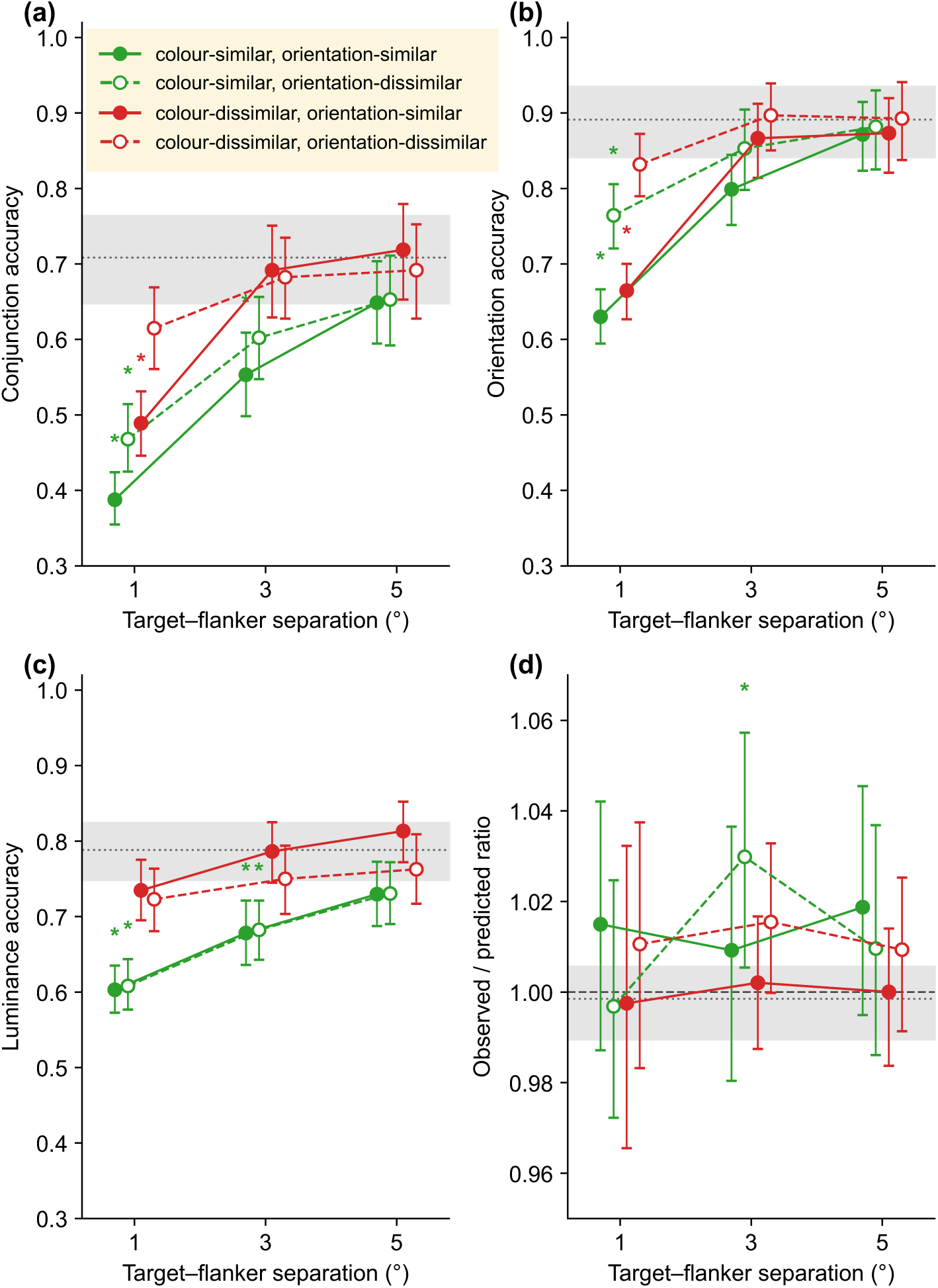
Accuracy on the 4AFC conjunction-report task as a function of target–flanker distance, flanker colour similarity to target, and flanker orientation similarity to target. Accuracy is the proportion of trials correct, plotted on the y-axis of each panel (a) – (c). (a) Conjunction accuracy: proportion of trials on which *both* target features were reported correctly. (b) Orientation accuracy: proportion correct for target orientation, irrespective of the luminance response. (c) Luminance accuracy: proportion correct for target luminance, irrespective of the orientation response. (d) Channel-independence ratio: observed conjunction accuracy divided by the value predicted if the two channels were independent (the product of the orientation and luminance accuracies in the same cell). A ratio of 1 (dashed horizontal line) indicates independence. Values above and below 1 indicate positive and negative coupling between channels. In (a)–(c), the horizontal dotted line and grey band show the target-alone reference (across-participant mean and 95% confidence interval), and asterisks mark cells whose 95% confidence interval falls entirely below this band. In (d), the dashed line marks a ratio of 1 (independence), and the dotted line and grey band show the target-alone ratio and its bootstrap 95% confidence interval, matching the reference shown in (a)–(c). Asterisks mark cells whose bootstrap 95% confidence interval excludes 1. Across all panels the four flanker conditions are coded by colour (green = colour-similar, red = colour-dissimilar) and by symbol (filled/solid = orientation-similar, open/dashed = orientation-dissimilar). Markers are offset horizontally within each distance for legibility.

### Conjunction accuracy

We first examined whether participants correctly reported both target features on each trial (Fig. 1a). A binomial GLMM with logit link revealed a robust main effect of target–flanker distance, χ²(2) = 406.18, *p* < .001, with conjunction accuracy increasing monotonically from 1° to 5°. Holm-corrected pairwise contrasts confirmed reliable improvement at every step (all *z* ≤ −4.72, all *p* < .001), indicating that crowding effects on the joint outcome extended somewhat beyond Bouma’s critical spacing (Bouma, 1970) in the conjunction task.

The two flanker-similarity dimensions had qualitatively distinct effects on conjunction accuracy. Colour-similar flankers reduced accuracy at all distances, χ²(1) = 148.43, *p* < .001, with comparable costs at 1° (Δ = −0.55 logit units, SE = 0.06, *z* = −8.91, Holm *p* < .001) and 3° (Δ = −0.51, SE = 0.06, *z* = −8.08, *p* < .001), and a smaller but still reliable cost at 5° (Δ = −0.27, SE = 0.07, *z* = −4.18, *p* < .001). The distance × colour interaction was reliable, χ²(2) = 11.06, *p* = .004. Orientation-similar flankers produced a stronger distance-dependent decrement: the main effect was reliable, χ²(1) = 19.32, *p* < .001, but the cost was confined to 1° (Δ = −0.46, SE = 0.06, *z* = −7.42, Holm *p* < .001), with no reliable effect at 3° (Δ = −0.09, SE = 0.06, *z* = −1.36, *p* = .351) or 5° (Δ = +0.06, SE = 0.07, *z* = +0.93, *p* = .354). This produced a significant distance × orientation interaction, χ²(2) = 36.07, *p* < .001. The colour × orientation interaction was not reliable, χ²(1) = 1.06, *p* = .304, indicating that the orientation-similarity cost did not depend on whether flankers also shared the target’s colour. The three-way interaction was significant, χ²(2) = 7.71, *p* = .021, of small magnitude and not interpreted further.

Comparison with the target-alone reference band identified four cells whose 95% CIs fell entirely below the lower edge of the target-alone band (Fig. 1a, asterisks): three of the four conditions at 1° (excluding only colour-dissimilar/orientation-dissimilar flankers, which sat at the lower edge of the target-alone 95% CI band), and the colour-similar/orientation-similar condition at 3°. All cells at 5° were within the target-alone band.

In summary, conjunction accuracy in the 4AFC task was shaped by two qualitatively distinct flanker effects. Colour-similar flankers produced a graded, spatially extensive decrement that remained reliable to 5°. Orientation-similar flankers produced a stronger decrement confined to the smallest target–flanker separation. To ask whether these two flanker dimensions act on separable report channels, we next examined accuracy on each feature considered alone.

### Orientation accuracy

We recoded each trial as orientation-correct if the participant’s orientation response matched the target’s orientation, regardless of whether the luminance response was also correct (Fig. 1b). The same binomial GLMM structure was applied to this binary outcome.

Orientation accuracy was influenced by all three factors. There was a main effect of target–flanker distance, χ²(2) = 430.96, *p* < .001, with accuracy improving as flankers moved further from the target (all Holm-corrected pairwise contrasts reliable, all *z* ≤ −4.72, all *p* < .001). Flanker orientation similarity produced a main effect, χ²(1) = 91.65, *p* < .001, that interacted with distance, χ²(2) = 43.76, *p* < .001. The interaction reflects the spatial extent of the orientation-similarity cost: orientation-similar (vertical) flankers impaired orientation reports substantially at 1° (Δ = −0.88 logit units, SE = 0.07, *z* = −12.28, Holm *p* < .001), less so at 3° (Δ = −0.37, SE = 0.09, *z* = −4.18, *p* < .001), and not at all at 5° (Δ = −0.15, SE = 0.09, *z* = −1.64, *p* = .101). This is the canonical Bouma profile: interference is confined to the critical-spacing window and is absent beyond it.

Flanker colour similarity also reduced orientation accuracy, χ²(1) = 36.19, *p* < .001, though the cost was smaller than that of orientation similarity. The colour-similarity cost was reliable at 1° (Δ = −0.32 logit units, SE = 0.07, *z* = −4.51, Holm *p* < .001) and 3° (Δ = −0.49, SE = 0.09, *z* = −5.58, *p* < .001) but had vanished by 5° (Δ = −0.07, SE = 0.09, *z* = −0.73, *p* = .468), producing a reliable distance × colour interaction, χ²(2) = 11.02, *p* = .004. The colour × orientation interaction was not significant, χ²(1) = 1.04, *p* = .307, nor was the three-way interaction, χ²(2) = 3.15, *p* = .207.

Three cells fell reliably below the target-alone reference band (Fig. 1b, asterisks): the three conditions at 1° in which flanker orientation matched the target. The colour-dissimilar/orientation-dissimilar condition at 1° sat just at the band’s lower edge. All cells at 3° and 5° were within the target-alone band.

In summary, the orientation-report channel was selectively impaired within Bouma’s window, with the impairment driven primarily by flanker orientation similarity and to a smaller extent by flanker colour similarity. This pattern matches the asymmetric dissociation we have reported previously in our orientation-only 2AFC task (Cass & Van der Burg, 2023), providing a within-task analogue of that finding.

### Luminance accuracy

The luminance-report channel showed an inverse pattern (Fig. 1c). Here the main effect of flanker colour similarity was reliable, χ²(1) = 144.42, *p* < .001. Colour-similar flankers reduced luminance accuracy reliably at every distance: at 1° by Δ = −0.59 logit units (SE = 0.06, *z* = −9.18, Holm *p* < .001); at 3° by Δ = −0.47 (SE = 0.07, *z* = −6.92, *p* < .001); and at 5° by Δ = −0.34 (SE = 0.07, *z* = −4.82, *p* < .001). The distance × colour interaction was reliable, χ²(2) = 6.98, *p* = .030: the colour-similarity cost on luminance reports was largest at 1° and shrank as flankers moved further from the target, though it remained reliable at every separation tested.

Flanker orientation similarity had a much smaller effect on luminance accuracy, χ²(1) = 5.48, *p* = .019, and within-distance contrasts revealed no reliable orientation-similarity cost at any separation: 1° (Δ = +0.02 logit units, SE = 0.06, *z* = +0.31, Holm *p* = .754), 3° (Δ = +0.10, SE = 0.07, *z* = +1.43, *p* = .305), or 5° (Δ = +0.16, SE = 0.07, *z* = +2.23, *p* = .078). The distance × orientation interaction was not significant, χ²(2) = 2.08, *p* = .353. The colour × orientation interaction was reliable, χ²(1) = 7.55, *p* = .006, with the colour-similarity cost being somewhat larger when flankers also differed from the target in orientation. The three-way interaction was not significant, χ²(2) = 1.61, *p* = .448.

Distance produced a substantial main effect, χ²(2) = 100.70, *p* < .001. The four colour-similar conditions at 1° and 3° fell reliably below the target-alone reference band (Fig. 1c, asterisks). Notably, the colour-similar cells remained substantially below the band at 3°, beyond Bouma’s window for this eccentricity, while the colour-dissimilar cells returned to baseline by 3°.

In summary, the luminance-report channel was selectively impaired by flanker colour similarity, with a spatially extensive decrement that remained reliable at 3° and 5°. Flanker orientation similarity had no reliable cost on luminance accuracy at any distance. This is the inverse pattern to the orientation-report channel and matches the asymmetric dissociation reported in our previous luminance-only 2AFC task (Cass & Van der Burg, 2023).

### Channel independence

The channel-specific analyses establish that the two report channels have distinct flanker sensitivities. They do not, however, establish that the two channels operate statistically independently within a given trial: orientation and luminance responses could in principle be associated at the trial level even when their flanker sensitivities differ. We tested for such trial-level coupling with two complementary analyses. The first, a ratio test (Fig. 1d), compares the observed conjunction accuracy in each cell of the design to the value that would be expected if the two channels were producing their reports independently (the product of the two marginal accuracies). The second analysis (Fig. 2) examines the full 4×4 stimulus-response matrix for each cell of the design. This matrix tabulates how often each of the four possible responses (left-bright, right-bright, left-dim, right-dim) is given to each of the four target identities. For each of the four stimulus rows of the matrix, we ask whether the four response probabilities are what would be expected if the orientation and luminance reports were generated by two independent decisions. The two tests probe complementary aspects of the joint response distribution and are sensitive to different ways in which the two channels could be coupled, as we elaborate when reporting each.

**Figure 2.**
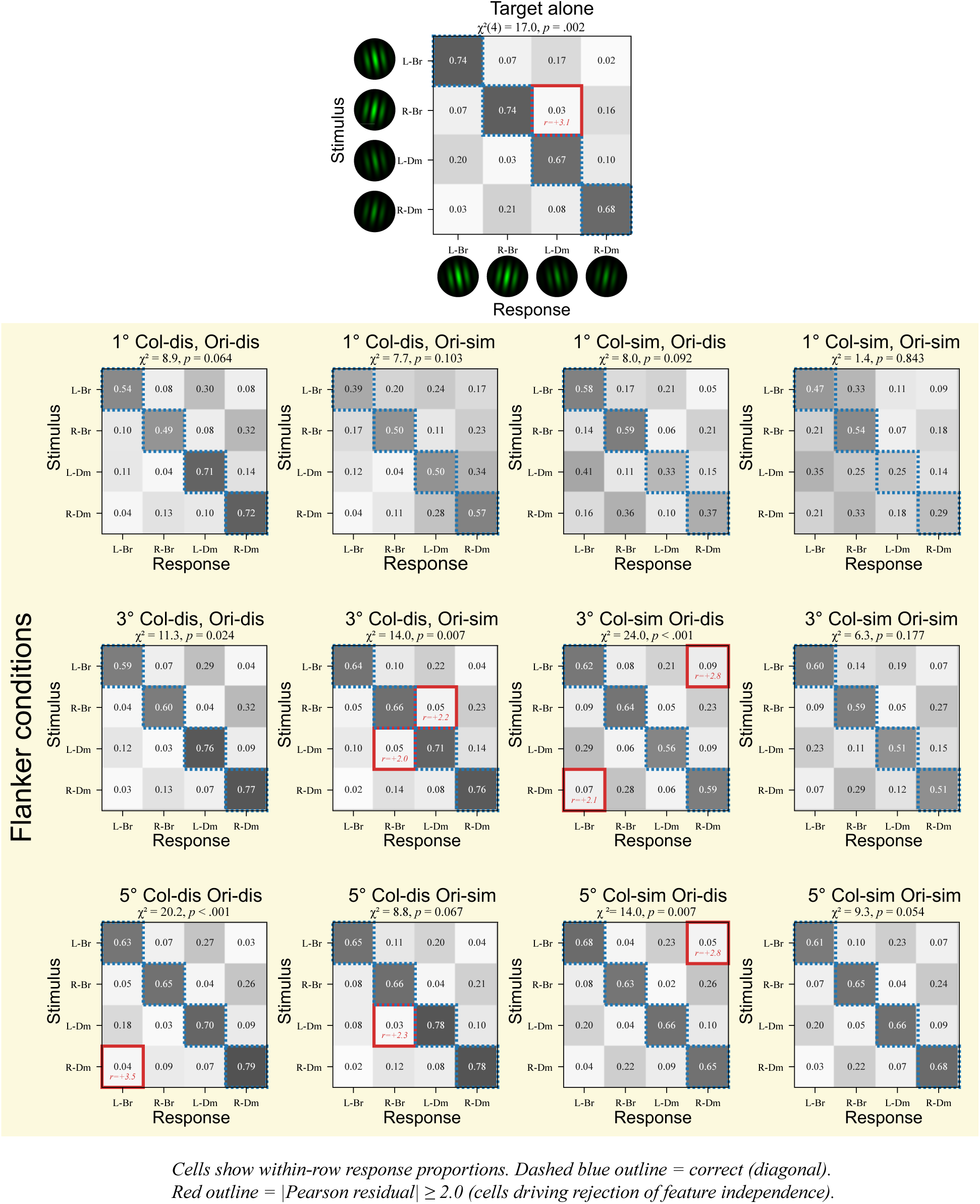
Stimulus-response matrices for the thirteen conditions of the design (twelve flanker conditions, yellow shading, plus the target-alone baseline). Each panel shows a 4×4 matrix in which rows are target identities (L-Br = left-tilt bright, R-Br = right-tilt bright, L-Dm = left-tilt dim, R-Dm = right-tilt dim) and columns are response identities, in the same order. Each cell shows the proportion of trials on which the row stimulus elicited the column response, normalised within rows; cell shading scales with this proportion. Diagonal cells (correct conjunction responses) are outlined in dashed blue. Cells with an absolute Pearson residual ≥ 2.0 against the within-row feature-independence expectation are outlined in red and labelled with the residual value, identifying the cells that drive any chi-square rejection in that panel. The χ² statistic and *p* value for each panel test of feature independence are reported in the panel title (df = 4 for all panels). Residuals are concentrated in specific off-diagonal entries rather than distributed uniformly, and the strongest residuals appear at 3° and 5° target–flanker separations rather than in the most heavily crowded 1° cells.

### Ratio test

For each of the twelve flanked cells, we computed the value of joint accuracy that would be expected under statistical independence of the two channels:

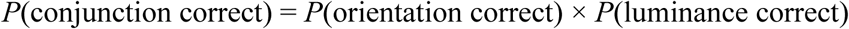

with both marginal probabilities computed as observed proportions correct, within participant and cell, rather than from the GLMM estimated marginal means. The ratio of observed to predicted joint accuracy provides a quantitative index of coupling between the two report channels. A ratio of one indicates independence: the two features are correct together exactly as often as their separate accuracies predict. A ratio above one indicates positive coupling, meaning the two channels tend to succeed together, and to fail together, more often than independence predicts, which is the pattern expected if crowding corrupts a bound representation of both features at once. A ratio below one indicates negative coupling, meaning the two errors are anti-associated, so that a trial on which one feature is misreported is one on which the other is more likely than chance to be reported correctly.

For each participant and cell we formed the ratio of observed joint accuracy to the product of the two marginal accuracies, and averaged these per-participant ratios across the sample. We obtained 95% confidence intervals by resampling participants with replacement (5,000 bootstrap iterations), recomputing the mean ratio on each resample, and taking the 2.5th and 97.5th percentiles. Because each participant contributes a single ratio, this procedure tests independence within observers and is not distorted by between-participant differences in overall accuracy. Bootstrap distributions for all cells are reported in Fig. 1d.

Across the twelve flanked cells, the observed-to-predicted ratio sat close to unity, ranging from 1.00 to 1.03 with no systematic departure in either direction. In eleven of the twelve cells, the bootstrap confidence interval on the ratio included unity, so observed joint accuracy was statistically indistinguishable from the product of the two single-feature accuracies. The single exception (colour-similar, orientation-dissimilar at 3°: ratio = 1.03, 95% CI [1.01, 1.06]) departed only marginally, was not the most heavily crowded condition, and was one reliable departure in twelve tests. Critically, the most heavily crowded cell (1° separation, colour-similar, orientation-similar) was fully consistent with independence (ratio = 1.02, 95% CI [0.99, 1.04]), so the small residual departures did not grow with crowding and were not largest where crowding was strongest. Across the full range of crowding strengths tested, conjunction accuracy is well approximated by the product of the two marginal accuracies. At the target-alone baseline the ratio was likewise consistent with independence (1.00, 95% CI [0.99, 1.01]).

### Chi-square tests of stimulus-response matrices

The ratio test above asks a single summary question: does the chance of getting both features right match the product of the two single-feature accuracies? It treats the three ways of being wrong as interchangeable. A second analysis lets us look inside the errors themselves. On every trial the observer chooses one of four responses, formed by crossing the two orientation answers (left, right) with the two brightness answers (bright, dim). We can therefore tabulate, for each of the four possible targets, how often it drew each of the four possible responses. We refer to this 4×4 table of stimulus against response as a stimulus-response matrix, and Figure 2 shows the matrix for each condition of the experiment.

The question we put to each table is simple: when the observer makes an error, are the two features corrupted separately, or together? Consider a bright left-tilted target. If the two judgements are made independently, then whether the observer gets the orientation right should tell us nothing about whether they get the brightness right. Errors would then scatter across the wrong responses in proportion to how often each feature is misjudged on its own. If instead the two judgements are linked, the errors would cluster: for example, getting the orientation wrong would go hand in hand with getting the brightness wrong, so that the observer lands on the response that is wrong about both. A chi-square test is a standard way of measuring this. It compares the counts actually observed in the table against the counts expected if the two features were judged independently, and returns a single number that grows as the observed table departs from that independent prediction. A small chi-square means the errors behave as two separate feature judgements; a large one means the two judgements are yoked together in at least some part of the table.

We computed this test for all thirteen conditions in our design: the twelve flanker conditions (three separations crossed with the two colour and two orientation flanker types) plus the target-alone baseline. For each condition we obtained one chi-square value, testing whether that condition’s 4×4 table is consistent with the two features being judged independently. A statistically significant value means the errors in that condition are not fully independent: orientation mistakes and brightness mistakes tend to occur on the same trials more (or less) often than chance. A non-significant value means the errors are well described by two separate feature judgements.

The test reached significance in six of the thirteen conditions (Table 1). The key point is where these six fell. Only one of them was a strongly crowded condition. These significant departures were concentrated at the intermediate and far separations (3° and 5°), and were absent at the most crowded 1° separation, where crowding is strongest. One of the six was the target-alone baseline, in which no flankers were present at all. If crowding worked by fusing the two features into a single object whose identity is then misread, the coupling between the two feature errors should have been strongest exactly where crowding is strongest. We observed the opposite pattern, which points away from a crowding-driven binding failure and toward a small coupling that is present regardless of crowding. We take up its likely source in the Discussion.

**Table 1.**
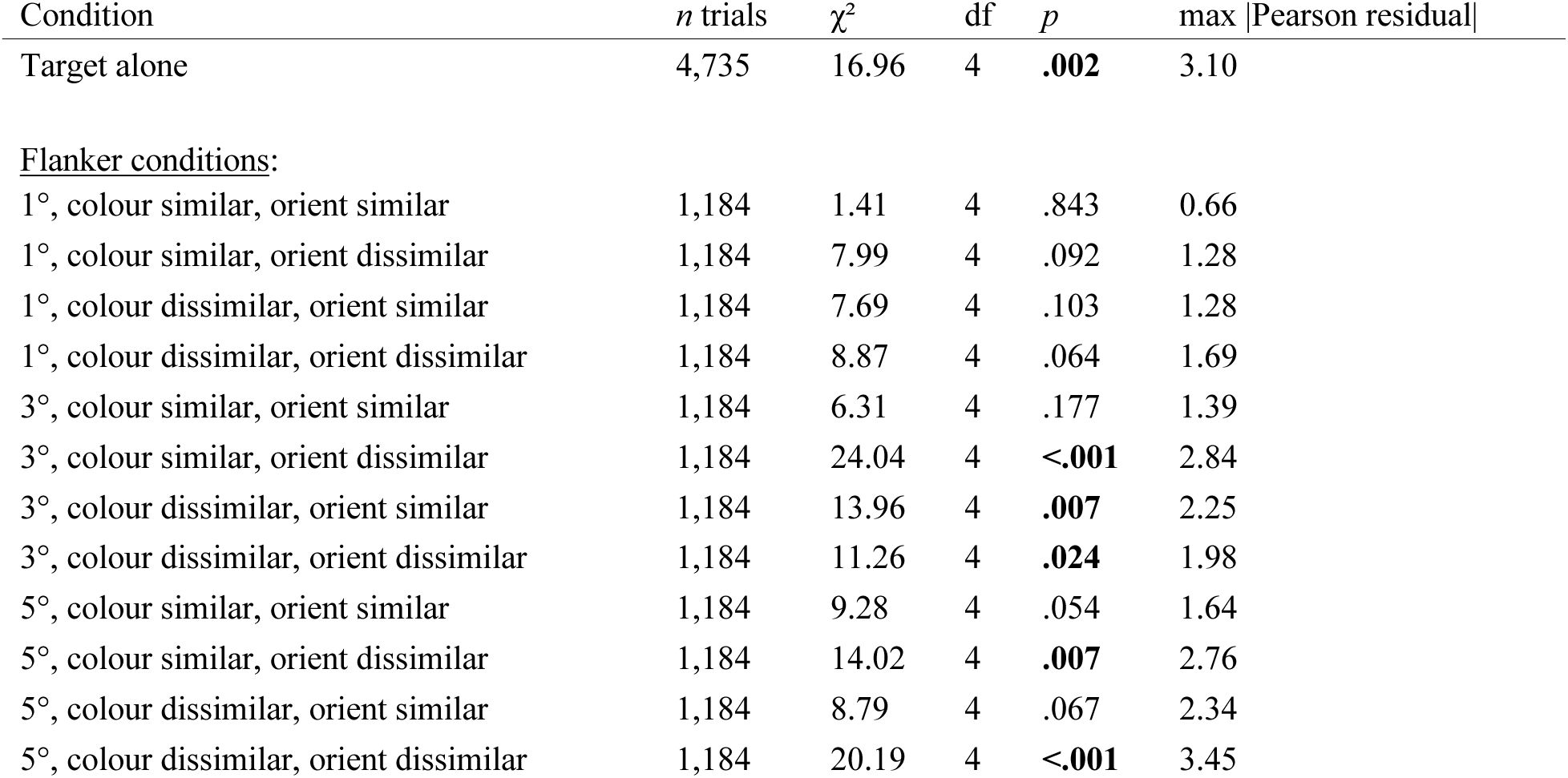
Chi-square test of within-stimulus feature independence in the 4×4 stimulus-response matrix, by design cell.

A rejection at the target-alone condition is informative because no flankers were present, so the rejection cannot reflect crowding-induced binding failure. We inspected the per-row Pearson residuals (red-outlined cells in Fig. 2) to see which response patterns drove the rejection. For the target-alone matrix, when participants reported the wrong orientation for a bright target they were disproportionately likely to also report dim luminance (Pearson residual = 3.10). In other words, a small fraction of trials produced an “opposite-corner” response in which both features were misreported in the same direction. This is the signature of a residual response bias rather than of feature binding, and is consistent with the bias structure documented in our earlier study ((Cass & Van der Burg, 2023), Fig. 5). The same kind of structure was visible in the flanked conditions in which the chi-square rejected independence: the over-represented cells were specific off-diagonal entries, not concentrated on the “both features wrong” opposite-corner cell that a bound-feature account would predict.

The two tests therefore deliver complementary verdicts. The ratio test confirms that observed joint accuracy is well predicted by the product of the marginal accuracies, with eleven of twelve cells statistically consistent with independence and no departure that grows with crowding or peaks in the most heavily crowded cell. The chi-square test refines this picture, showing that the small residual structure in the off-diagonal response cells is present even at target-alone and does not track crowding strength. We consider the implications of this combined pattern for the bound-features versus independent-features accounts of crowding in the Discussion.

## Discussion

We asked whether crowding of a two-feature target reflects the disruption of a bound object representation, or whether apparent object-level failures can be reconstructed from largely separable distortions of the target’s component features. To address this we employed a 4AFC conjunction-report task in which observers reported both the brightness and the orientation of every target on every trial, so that each feature judgement was made on the very same target, viewed under the same conditions. The results favoured the latter account. Conjunction accuracy declined robustly under crowding, but the structure of these errors was not unitary. Orientation reports were primarily impaired by orientation-similar flankers within the smallest target–flanker separations, whereas luminance reports were primarily impaired by target-colour-similar flankers across a broader spatial range. Critically, the joint accuracy actually observed was closely predicted by simply multiplying the two single-feature accuracies (Figure 1d). This is the prediction one would make if the two report channels failed independently, and it held in eleven of the twelve flanked conditions and at the target-alone baseline, with observed joint accuracy statistically indistinguishable from the product of the marginal accuracies. A single condition departed marginally from independence, and it was neither the most heavily crowded condition nor part of any orderly pattern across separations. A genuine binding failure predicts the opposite: coupling that emerges and grows where crowding is strongest, in the most heavily crowded condition of all (similar colour and similar orientation at the smallest separation). That condition was fully consistent with independence. The coupling that a binding failure would produce was therefore absent exactly where that account requires it to be strongest. On many crowded trials observers reported the conjunction incorrectly, but these errors arose predominantly from independent failures of the two feature reports rather than from disruption of a representation in which the two features are bound.

This result builds on a pattern of findings showing that object-level crowding errors can have heterogeneous origins. Dakin et al. (2010) studied this with letter-like T stimuli abutted by single flankers, and found that the structure of the errors depended on where the flanker was placed in object-centred coordinates. When the flanker was located to the side of the T (regardless of the T’s retinal orientation), errors were well captured by positional averaging of the component features. When the flanker sat at the end of the T, errors instead resembled substitutions of the flanker for the target: a different kind of object-level error in which the whole flanker effectively replaces the target. The present data extend the feature-level account to independently varied feature dimensions. Orientation and luminance fail under crowding at rates that combine multiplicatively, just as two independent channels would predict. This is a third route by which crowding produces object-level errors. Three different mechanisms can therefore produce the same outcome on a conjunction-report task: feature-level positional averaging of mislocalised features (Dakin et al.’s side-flanker condition); object-level substitution of the whole flanker for the target (Dakin et al.’s end-flanker condition); or two independent feature-channel failures that happen to co-occur on a trial (the present results). Accuracy alone does not tell these three apart, because each produces incorrect conjunction reports. The full stimulus-response matrix does, because each mechanism leaves a different pattern in the joint distribution of errors across the four response identities. Positional averaging concentrates errors on the response that interpolates the target and flanker features; whole-object substitution concentrates them on the response matching the flanker; independent feature-channel failures distribute them according to the product of the two marginal error rates. Which mechanism is operating in a given task depends on the stimulus configuration and the feature pair tested. The stimulus-response matrix provides a means for distinguishing them.

We are careful not to over-claim from this. The present data do not show that feature binding is never vulnerable to crowding. The broader literature suggests the answer depends on which feature pair is interrogated: orientation and spatial frequency show compulsorily bound errors under crowding (Yashar et al., 2019), as do orientation and position (Greenwood et al., 2009), while hue and motion direction (Greenwood & Parsons, 2020) and hue and orientation (Yashar et al., 2019) behave more like the orientation × luminance pair tested here. Greenwood and Parsons (2020) in particular used a within-trial joint-report design closely analogous to ours, and observed that crowding strength could be manipulated independently for each feature dimension, with errors dissociating along the manipulated channel. The present results extend this conjunction-report logic to a new feature pair (orientation × luminance) and add a complementary analytic test (the ratio test plus chi-square) for the same channel-independence claim. What the present data establish is the more modest claim that, when orientation and luminance are varied orthogonally in a conjunction-report task, the observed conjunction cost does not exceed what would be predicted from independent feature-level failures.

The two channels also differed in spatial signature. The orientation-similarity cost on orientation accuracy was confined to the smallest separation, matching the canonical Bouma profile (Bouma, 1970). The colour-similarity cost on luminance accuracy was broader, remaining reliable at 5°, beyond Bouma’s window for our 6° eccentricity. This spatial-signature dissociation suggests that the integration zones of the two channels are not feature-general, and is consistent with the suggestion that hue-contingent luminance crowding engages mechanisms with larger integration zones than the orientation regime, perhaps reflecting a hue-grouping or surface-formation stage distinct from classical contour-integration crowding (Cass & Van der Burg, 2023).

A final question is whether the feature-independence we report would also hold in dense displays. The present experiment used sparse displays, a single target flanked by two Gabors, for which Bouma’s law provides a good account of the spatial extent of crowding. In dense displays Bouma’s law does not hold. Instead, what determines crowding is the featural similarity of the target’s immediately adjacent neighbours, which can lie much closer to the target than Bouma’s window would predict (Van der Burg et al., 2024; Van der Burg, 2017). The CODE model formalises this by allowing the neural response to each element to be influenced by its nearest neighbours according to their proximity and similarity, and accounts for crowding in both sparse and dense displays within a single framework (Van der Burg et al., 2024). Dissimilar nearest neighbours release the target from crowding even when similar, otherwise-deleterious flankers remain within Bouma’s window, so that crowding is set by the immediately adjacent elements rather than by the full contents of the critical-spacing window (Van der Burg, 2017), a pattern with which recent findings are broadly consistent (Chakravarthi, 2026). Whether the two feature channels we find to be independent in sparse displays remain independent in dense ones is an open question, and the nearest-neighbour framework reframes it. Because crowding in dense displays is set by the similarity of the nearest neighbours, the relevant manipulation is not the overall density of the array but the featural relation of the few elements immediately flanking the target. Our sparse results predict that when those nearest neighbours share only one feature dimension with the target (only colour, or only orientation), that dimension should be crowded while the other is comparatively spared, preserving channel independence. If instead the nearest neighbours are similar to the target on both dimensions, the two channels might fail together and yield a genuinely object-level cost that is absent in the sparse case. Testing this requires holding the immediate neighbourhood of the target under experimental control while varying its featural similarity, together with the conjunction-report measure used here, and is a natural next step for distinguishing feature-level from configural crowding.

A further question is whether the channel independence we observe extends to feature pairs other than orientation and brightness. The wider evidence is mixed. Some pairs dissociate under crowding, implying separable channels: hue and orientation are reported independently (Yashar et al., 2019), as are colour and motion direction (Greenwood & Parsons, 2020). Others instead produce correlated errors of the kind expected when two features are bound rather than judged separately, as for orientation and spatial frequency (Yashar et al., 2019) and for relative position and orientation (Greenwood et al., 2012). The evidence for a given pair can also depend on what is measured, since positional averaging of component features can itself generate apparently object-level errors without a dedicated binding stage (Dakin et al., 2010). The two descriptions coincide when the two features are averaged together rather than separately (e.g. one at a time). One organising possibility, advanced by Yashar et al. (2019), is that binding tracks shared neural coding: the pairs that bind are jointly encoded within single neurons in early visual cortex, which are tuned to orientation, spatial frequency, and position together (Hubel & Wiesel, 1968) (De Valois et al., 1982), whereas the pairs that dissociate are carried by more segregated pathways. According to this view, two features crowd as a unit when a common population encodes both, and independently when separate populations encode each.

Our conjunction-report framework offers a direct way to test this. Because both features are reported on every trial for the same target under identical conditions, the joint distribution of the two responses is available directly, rather than being inferred by comparing separate blocked tasks. The ratio test then indexes the strength of coupling (unity under independence), while the chi-square analysis of the full stimulus-response matrix is sensitive to the particular response confusions a binding failure produces. Applying both across a parametric manipulation of target– flanker similarity and separation is especially diagnostic, because genuine binding predicts coupling that grows with crowding strength, whereas a similarity-independent response bias predicts coupling that is present even without flankers. Mapping feature pairs in this way offers a principled approach to specifying which features are bound under crowding and which remain independent.

### Reconciling the ratio test and the chi-square analysis

The two tests ask different questions of the same data. The ratio test asks whether observers get both features right as often as their separate accuracies predict. The answer is yes in eleven of the twelve flanker conditions, and the single exception falls at an intermediate separation rather than the most crowded one. The chi-square test asks a more demanding question, and rejects independence in five conditions. Both tests agree on the main point, that crowding leaves no binding signature, but the chi-square picks up structure in the errors that the ratio test does not.

The difference lies in what each test pools over. The ratio test collapses across the four possible targets and asks a single question: across the experiment as a whole does getting the orientation right tell us anything about getting the brightness right? The chi-square asks that question separately for each of the four targets. For a bright left-tilted target it asks whether the observer’s orientation and brightness responses are independent on those trials alone, then repeats this for the other three targets and combines the results. Because it never averages across targets, the chi-square can detect structure that averaging would wash out.

That structure is real, though in most cells it is very slight. It falls consistently in one place: *the response that is wrong about both features at once*. Only a handful of cells show a departure large enough to reach significance on its own (Figure 2, red outlines), but the direction is highly consistent. Across the thirteen conditions and four targets, the *both-wrong* response was more common than independence predicts in forty-five of the fifty-two cells, where chance predicts twenty-six (mean Pearson residual = +0.90, *t*(51) = 6.34, *p* < .001). Individually these departures are small, amounting to roughly one extra trial in a hundred. Collectively they reveal a systematic tendency for observers who err to err about both features together.

We can be clear about three things this tendency is not. It is not a binding failure produced by crowding. A bound representation corrupted by flankers would yield a coupling that emerges under crowding and strengthens as crowding intensifies. Instead, the coupling is present in full at the target-alone baseline, where no flankers were shown and crowding cannot apply, and it does not grow when flankers are closer to the target. A tendency already present without flankers, and unchanged by them, cannot be a product of crowding. It is also not a simple guessing artefact.

Guessing offers a simple explanation: because a single key reports both features, a wrong guess is often wrong about both. If that were the cause, the coupling should be strongest where accuracy is lowest and guessing should be most frequent. Accuracy and the size of the coupling were not related in this way (r = 0.43, *p* = .17), and in the opposite direction to the guessing prediction. Finally, it is not the directional response bias reported by Cass and Van der Burg (2023), who found a tendency to report targets as bright under green vertical flankers. That bias concerns which response an observer favours within a single feature, and the blocked design used there, in which brightness and orientation were judged in separate trials, cannot produce coupling between two features at all. The present coupling is a relation between two judgements made on the same trial, and only a conjunction task can reveal it. We cannot yet establish is why the response coupling occurs, or why it varies across conditions as it does. We leave this as a question for further work. For present purposes, the important point is that whatever generates this small response coupling, it operates independently of crowding.

## Conclusion

We set out to ask a basic question about how the visual system fails under conditions which produce weak and strong crowding. Does crowding disrupt the recognition of an object as a unified whole, compromising the joint representation of the features that belong to it, or does it instead degrade the recognition of the object’s component features separately, so that the object-level confusions we observe simply fall out of two separate feature failures? We addressed this with a 4AFC conjunction-report task that required observers to report both the orientation and the brightness of every target on every trial, analysing the resulting performance from three angles: at the level of joint accuracy, at the level of each feature considered alone, and at the level of statistical coupling between the two featural report channels.

The picture that emerged was consistent across all three analyses. Orientation reports and luminance reports were impaired by flankers in qualitatively different ways and across different spatial ranges, indicating that the two channels are tuned to different similarity dimensions and have different integration zones. When we asked whether the two channels’ errors were coupled at the trial level, the answer was that they were not: observed joint accuracy was closely predicted by the product of the two marginal accuracies in every cell of the design, and the small departures from this prediction did not concentrate at the most heavily crowded condition. The conjunction errors observed under crowding were constituted from two largely independent feature-specific failures, rather than from disruption of a representation in which the two features are bound.

For object recognition more broadly, this matters because it speaks to how peripheral clutter limits perception. A visual system in which crowding compromises bound object representations would be a one in which information about cluttered objects is lost wholesale. The present results suggest a more selective kind of failure: when one feature dimension of an object is compromised by clutter, the other can remain partially accessible, and the object’s identity may be partly reconstructed from whichever channel happens to survive the local interference. That this pattern coexists with the *opposite* pattern for other feature pairs in the literature (orientation and spatial frequency, orientation and position) suggests that the architecture of crowding is feature-pair-specific rather than uniform across the visual system. Some features are bound at encoding and fail together, whilst others remain separable and fail in their own ways. Mapping which feature pairs follow which pattern, and why, is the natural next step.

## Declaration of generative AI and AI-assisted technologies in the manuscript preparation process

During the preparation of this work, the authors used Claude (Anthropic) to assist with drafting and editing the text of the manuscript. The authors reviewed and edited the output as needed and take full responsibility for the content of the published article.

